# A structured evaluation of cryopreservation in generating single cell transcriptomes from cerebrospinal fluid

**DOI:** 10.1101/2021.09.15.460354

**Authors:** Hanane Touil, Tina Roostaei, Daniela Calini, Claudiu Diaconu, Samantha Epstein, Catarina Raposo, Licinio Craveiro, Ilaria Callegiri, Julien Bryois, Claire S. Riley, Vilas Menon, Tobias Derfuss, Philip L. De Jager, Dheeraj Malhotra

**Author notes:** contributed equally (Hanane Touil, Tina Roostaei, Daniela Calini).

## Abstract

**Importance:** A robust cerebrospinal fluid (CSF) cell cryopreservation protocol using high resolution single-cell (sc) transcriptomic data would enable the deployment of this important modality in multi-center translational research studies and clinical trials in which many sites do not have the expertise or resources to produce data from fresh samples. It would also serve to reduce technical variability in larger projects.

**Objective:** To test a reliable cryopreservation protocol adapted for CSF cells, facilitating the characterization of these rare, fragile cells in moderate to large scale studies.

**Design:** Diagnostic lumbar punctures were performed on twenty-one patients at two independent sites. Excess CSF was collected and cells were isolated. Each cell sample was split into two fractions for single cell analysis using one of two possible chemistries: 3’ sc-RNA-Sequencing or 5’sc-RNA-Sequencing. One cell fraction was processed fresh while the second sample was cryopreserved and profiled at a later time after thawing.

**Setting:** The research protocol was deployed at two academic medical centers taking care of multiple sclerosis and other neurological conditions.

**Participants:** 21 subjects (age 24 – 72) were recruited from individuals undergoing a diagnostic lumbar puncture for suspected neuroinflammatory disease or another neurologic illness; they donated excess CSF.

**Findings:** Our comparison of fresh and cryopreserved data from the same individuals demonstrates highly efficient recovery of all known CSF cell types. The proportion of all cell types was similar between the fresh and the cryopreserved cells processed, and RNA expression was not significantly different. Results were comparable at both performance sites, and with different single cell sequencing chemistries. Cryopreservation also did not affect recovery of T and B cell clonotype diversity.

**Conclusion and relevance:** Our cryopreservation protocol for CSF-cells provides an important alternative to fresh processing of fragile CSF cells: cryopreservation enables the involvement of sites with limited capacity for experimental manipulation and reduces technical variation by enabling batch processing and pooling of samples.

**Key points:** *Question:* How efficient is CSF cryopreservation for single-cell transcriptome analysis and can it be implemented in large multi-center translational and clinical trial settings?

*Findings:* We compared single-cell transcriptomes of paired fresh and cryopreserved CSF from 21 patients at two independent sites. We validate the efficacy of a simple and cost effective CSF cryopreservation method that preserves the composition and the transcriptomes of CSF cells stored for weeks-months. The protocol is deployed in a large multicenter Phase 4 MS clinical trial.

*Meaning:* A validated CSF cryopreservation method that would significantly advance basic science and biomarker research in neurological disorders by implementing single-cell transcriptome analyses in multi-center research and clinical trials.

## Introduction

The composition of cerebrospinal fluid (CSF) provides insights into the physiological and pathological states of the central nervous system (CNS). Analysis of CSF represents an important element in the diagnosis and monitoring of neurological diseases; it can also play a critical role in clinical trials to demonstrate target engagement and uncover adverse events. The fluid component of the CSF is sampled routinely in the diagnosis of CNS infections and a variety of neurological conditions for different biomarkers, including multiple sclerosis (MS) [oligoclonal bands]^(1)^, Alzheimer disease (AD) [Aβ/Tau]^(2)^, neuromyelitis optica (NMO) [AQP4 Ab]^(3)^, autoimmune encephalitis [anti-GAD and anti-NMDA-R antibody]^(4, 5)^. While proteins from the CSF supernatant are utilized for diagnostic purposes, currently the evaluation of the CSF cellular component is often limited to measuring cell composition at low resolution (measuring the presence of different major immune cell subsets) given their fragile nature and low numbers in CSF. Cytology for malignant cells is performed clinically when malignancies are considered. Research assays typically require rigorous sample handling and larger CSF volumes; using flow cytometry^(6)^, they have returned evidence of association between certain epitopes or cell subtypes and clinical outcomes. High resolution single cell analysis of fresh CSF samples has provided new insights into disorders such as MS^(7)^ and brain metastasis^(8)^.

Reliable methods to cryopreserve CSF-cells would significantly advance basic and translational research of neurological diseases. Here, we report an efficient protocol to cryopreserve CSF-cells for long-term storage, illustrating its utility using high resolution single-cell analysis. We show that the protocol is robust to batch effects, storage-time restrictions and sequencing chemistries. The protocol has now been deployed in a large multicenter Phase 4 MS clinical trial https://clinicaltrials.gov/ct2/show/NCT03523858?term=Consonance&draw=2&rank=1. We provide a detailed protocol and video with step-by-step instructions as a valuable resource for the community for broader implementation in clinical and translational research.

## Methods

### Study design

Diagnostic lumbar punctures (LPs) were performed on twenty-one patients who presented with suspected neuroinflammatory disorders to the Multiple Sclerosis Center at either Columbia University Irving Medical Center or University Hospital Basel. Six patients recruited at the Columbia University Medical Center and seven patients at the University Hospital Basel were analyzed using droplet-based 3’ single-cell RNA sequencing (3’ sc-RNA-Seq) on the 10x Genomics Chromium platform. Eight patients recruited at the University Hospital Basel were analyzed using 5’ sc-RNA-seq on the same platform. All participants provided written informed consent forms as part of a protocol approved by Columbia and Basel university’s institutional review board (IRB). All LPs were performed in sterile conditions. CSF samples were collected in sterile tubes, using the Sprotte spinal needle and immediately transported to the laboratory in an ice bucket to maintain a stable temperature of 0°C to 4°C. The volume and cell-count of CSF samples varied between 3-18mL, 615-5899 cells, respectively. The CSF samples were immediately processed upon arrival to the laboratory without delay. The number of donors and cells used for the analysis is described in (Table 1).

**Table 1:**
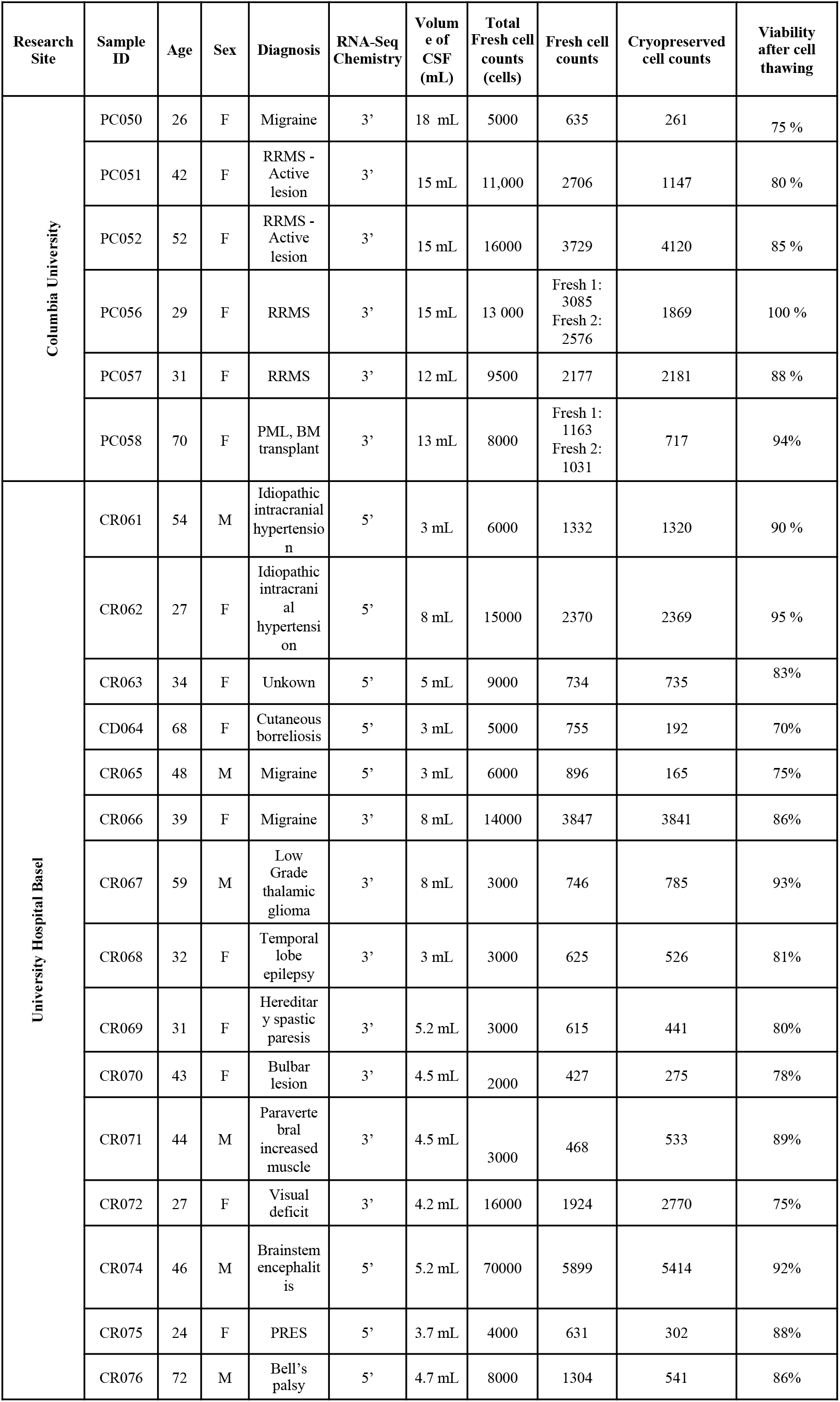
Patient demographics and sample description. • Estimated CSF-cell counts of freshly collected CSF samples before splitting into fresh and cryopreserved aliquots. • Fresh cell counts and cryopreserved cell counts represent cells called by CellRanger prior to the QC metrics sample/cell exclusion. • Viability of CSF-cell after thawing of cryopreserved cells is estimated using the automated cell counter.

| Research Site | Sample ID | Age | Sex | Diagnosis | RNA-Seq Chemistry | Volume of CSF (mL) | Total Fresh cell counts (cells) | Fresh cell counts | Cryopreserved cell counts | Viability after cell thawing |
| --- | --- | --- | --- | --- | --- | --- | --- | --- | --- | --- |
| Columbia University | PC050 | 26 | F | Migraine | 3' | 18 mL | 5000 | 635 | 261 | 75 % |
|  | PC051 | 42 | F | RRMS - Active lesion | 3' | 15 mL | 11,000 | 2706 | 1147 | 80 % |
|  | PC052 | 52 | F | RRMS - Active lesion | 3' | 15 mL | 16000 | 3729 | 4120 | 85 % |
|  | PC056 | 29 | F | RRMS | 3' | 15 mL | 13 000 | Fresh 1: 3085<br>Fresh 2: 2576 | 1869 | 100 % |
|  | PC057 | 31 | F | RRMS | 3' | 12 mL | 9500 | 2177 | 2181 | 88 % |
|  | PC058 | 70 | F | PML, BM transplant | 3' | 13 mL | 8000 | Fresh 1: 1163<br>Fresh 2: 1031 | 717 | 94% |
| University Hospital Basel | CR061 | 54 | M | Idiopathic intracranial hypertension | 5' | 3 mL | 6000 | 1332 | 1320 | 90 % |
|  | CR062 | 27 | F | Idiopathic intracranial hypertension | 5' | 8 mL | 15000 | 2370 | 2369 | 95 % |
|  | CR063 | 34 | F | Unkown | 5' | 5 mL | 9000 | 734 | 735 | 83% |
|  | CD064 | 68 | F | Cutaneous borreliosis | 5' | 3 mL | 5000 | 755 | 192 | 70% |
|  | CR065 | 48 | M | Migraine | 5' | 3 mL | 6000 | 896 | 165 | 75% |
|  | CR066 | 39 | F | Migraine | 3' | 8 mL | 14000 | 3847 | 3841 | 86% |
|  | CR067 | 59 | M | Low Grade thalamic glioma | 3' | 8 mL | 3000 | 746 | 785 | 93% |
|  | CR068 | 32 | F | Temporal lobe epilepsy | 3' | 3 mL | 3000 | 625 | 526 | 81% |
|  | CR069 | 31 | F | Hereditary spastic paresis | 3' | 5.2 mL | 3000 | 615 | 441 | 80% |
|  | CR070 | 43 | F | Bulbar lesion | 3' | 4.5 mL | 2000 | 427 | 275 | 78% |
|  | CR071 | 44 | M | Paravertebral increased muscle | 3' | 4.5 mL | 3000 | 468 | 533 | 89% |
|  | CR072 | 27 | F | Visual deficit | 3' | 4.2 mL | 16000 | 1924 | 2770 | 75% |
|  | CR074 | 46 | M | Brainstem encephalitis | 5' | 5.2 mL | 70000 | 5899 | 5414 | 92% |
|  | CR075 | 24 | F | PRES | 5' | 3.7 mL | 4000 | 631 | 302 | 88% |
|  | CR076 | 72 | M | Bell's palsy | 5' | 4.7 mL | 8000 | 1304 | 541 | 86% |

### CSF cell cryopreservation

CSF samples were centrifuged at 300g for 10 minutes at 4°C and the supernatant was aliquoted into 1mL aliquots and stored in the -80°C freezers. The CSF-cell pellet was resuspended in 70µL of CSF, and cell viability and counts were assessed using automated cell counters at Nexcelom (Nexcelom, Bioscience) with acridine orange and propidium iodide (AOPI) viability dye (Nexcelom, Bioscience).

The CSF-cell suspension was split into two aliquots: the first aliquot was used to analyze the transcriptomic profile of fresh CSF-cells using single-cell RNA sequencing (3’ and 5’sc-RNA-seq), while the second CSF aliquot was immediately cryopreserved and stored in the liquid nitrogen (LN) for 7 days to 2 months. For samples PC056 and PC058, we prepared three CSF-cell aliquots: two fresh (technical replicates) and the third aliquot was cryopreserved for up to 2 months. For cryopreservation, the CSF-cell suspension was diluted into 750µL of RPMI-1640 supplemented with 40% FBS in a gentle manner. We further diluted the CSF-cell suspension by dropwise adding 750uL of freezing medium (RPMI-1640 supplemented with 40% FBS and 20% DMSO) pre-chilled at 4°C. The CSF sample was immediately placed inside a freezing container (Mister Frosty, Thermo Fisher) and stored in the -80 °C freezer overnight, before it was moved to the LN. After the storage period elapsed, the cryopreserved cells were retrieved from the LN, and thawed at 37°C for 1-2 minutes. The thawed cells were subsequently diluted 1:4 using pre-warmed RPMI-1640 supplemented with 10% FBS and centrifuged at 300g for 10 minutes at room temperature. The supernatant was discarded and the cells were resuspended in fresh 60µL RPMI-1640 media, counted and processed for 3’ or 5’-single cell RNA-seq as in (Figure 1). The sample processing procedure was videotaped and is shared as the video link in supplementary figures (Video). The detailed protocol for sc-RNA-sequencing of fresh and cryopreserved CSF-cells as implemented in Roche Consonance trial is available here: https://clinicaltrials.gov/ct2/show/NCT03523858?term=Consonance&draw=2&rank=1

**Figure 1:**
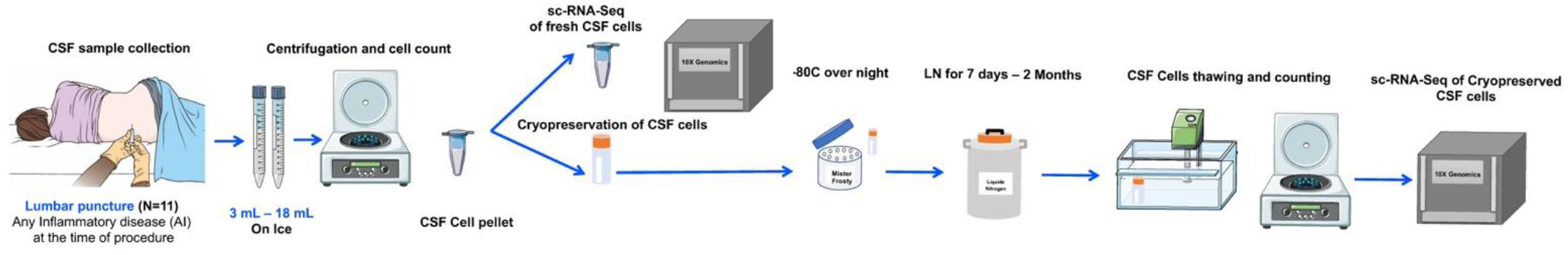
Schematic representation of the study design.

### Single-cell RNA and TCR-BCR sequencing

Fresh and cryopreserved CSF-cells from 44 libraries (14 from Columbia and 30 from Basel) were loaded into the 10x Genomics Chromium Controller for droplet-encapsulation. cDNA libraries were prepared using either the Chromium Next GEM Single Cell 3□ v3.1 or Chromium Next GEM Single Cell V(D)J v1.1 and v2.0 kits (10x Genomics) according to the manufacturer’s instructions. When the latter was used, TCR- and BCR-enriched libraries were prepared for each sample using Chromium Single Cell V(D)J Enrichment Kit (Human T Cell) and Chromium Single Cell V(D)J Enrichment Kit (Human B Cell) respectively. All libraries were sequenced using NovaSeq 6000 (Illumina) and NovaSeq 6000 S2 Reagent Kit v1.5 (100 cycles) (Illumina) to get a sequencing depth of 50K reads/cell (whole transcriptome libraries) or 10K reads/cell (TCR and BCR enriched libraries).

### Single-cell RNA data analysis

scRNA sequenced reads were aligned and quantified using Cell Ranger v3.1 to reference 3.1.0 (Ensembl 93) transcriptome for 3’ samples, and using Cell Ranger v6.0.2 multi pipeline to reference 2020-A (Ensembl 98) transcriptome and VDJ reference 5.0.0 (Ensembl 94) for 5’ whole transcriptome and TCR and BCR sequencing data.

The rest of the pre-processing and analysis were performed using Seurat library (v 4.0.1)^(10)^. For each sample, we performed quality control and filtered out cells: (1) that were likely doublets using Scrublet^(9)^, (2) with less than 100 genes, (3) with more than 25% mitochondrial transcripts, (4) Red Blood Cell (RBC) clusters (defined as clusters that showed high expression of hemoglobin genes). 3’ and 5’ samples were then independently merged, resulting in two separate datasets for subsequent analyses. The top 1000 variable genes were identified for each sample, aggregated over all samples in each dataset, and used for PCA dimensionality reduction. Integration was performed using the top 50 PCs using Harmony (https://doi.org/10.1038/s41592-019-0619-0). UMAP representation and clustering were performed using default Seurat parameters. Each cluster’s marker cells were identified using Seurat *FindAllMarkers* function for genes that were expressed in at least 25% of the cells from that cluster.

We annotated cells using a three-pronged approach to improve the annotation accuracy. (1) We performed differential gene expression (DE) of clusters using Seurat *FindAllMarkers* function to identify each cluster marker gene (gene expressed in at least 25% of the cells with fold change >1.28 at FDR <0.05 between clusters). (2) We also annotated clusters using a curated CITE-seq atlas reference (reference-based annotation) of human peripheral blood mononuclear cells, Azimuth^(12)^ developed as part of the NIH Human Biomolecular Atlas Project. Finally, we annotated cells by manual inspection of expression of marker genes from (1), (2), and published CSF single-cell RNA-seq studies.

We performed differential expression analyses between fresh and paired cryopreserved samples within each cluster using Seurat *FindMarkers* function with MAST^(13)^ test, fitting generalized linear models that are adapted for zero-inflated single-cell gene expression data. Patient ID was considered as a latent variable in order to account for the paired data structure. Genes detected in at least 10% of cells in either of the fresh or cryopreserved groups were included in the analyses. Groups with <5 cells were excluded from gene differential expression analysis. Genes with Bonferroni-adjusted p-value <0.05 and absolute log2 fold change >0.58 (fold change >1.5) were considered significant.

### Statistical analysis

Wilcoxon signed-rank test was used to compare recovered cell counts, RNA-sequencing quality control metrics, cluster cell frequencies, and frequencies of RBCs and low-quality cells between fresh and cryopreserved samples. Spearman’s ρ was used to assess correlation between variables. Reported p-values are all two-sided. Adjustment for multiple comparisons was performed using the Benjamini and Hochberg method. Enrichment of clonotypes with ≥2 cells in each cluster were assessed using one-sided Fisher’s exact test.

## Results

### Highly efficient cryopreservation of CSF-cells

We assessed the performance of our cryopreservation protocol by processing samples at two independent sites: six pairs of fresh and cryopreserved CSF samples using 3’ sc-RNA-Seq analysis at the Columbia University Irving Medical Center (CUIMC), USA, and fifteen pairs of fresh and cryopreserved CSF samples using both 3’ sc-RNA-Seq (N=7) and 5’ sc-RNA-seq (N=8) at University Hospital Basel (UHB), Switzerland.

#### 3’samples

Three out of thirteen fresh samples contained RBC clusters compared to none in the cryopreserved samples (p = 0.18). The proportion of low-quality cells excluded during QC (cells with <100 unique genes or a mitochondrial gene percentage >25%) were also not different between fresh and cryopreserved samples (p = 0.37) (Supplementary Table 1). After removing RBC, doublets and low-quality cells (details in the Methods), we retained 45,175 cells from 13 fresh-cryopreserved pairs and two additional fresh technical replicates for further analysis (Table 1). The number of QC passing cells was reduced but not significantly different between cryopreserved (median= 785 cells, IQR= 1,656) and fresh (median= 1,163 cells, IQR=2,088) samples (p = 0.24) (Figure 2A). Among sequencing QC metrics, the percentage of mitochondrial transcripts/cell was not significantly different in fresh (median= 2.6%, IQR= 1.6) versus cryopreserved (median= 3.0%, IQR= 2.1) (p = 0.68). The number of UMIs/cell in fresh (median= 4,396, IQR= 1,215) versus cryopreserved (median= 3,368, IQR= 2,851), and the number of unique genes/cell in fresh (median= 1,532, IQR= 279) versus cryopreserved (median= 1,199, IQR= 772) were reduced in cryopreserved samples (p = 0.01 and 0.008 for log-transformed values, respectively), but remained within the acceptable range of parameters for downstream analysis (Supplementary Figure 1A).

**Figure 2:**
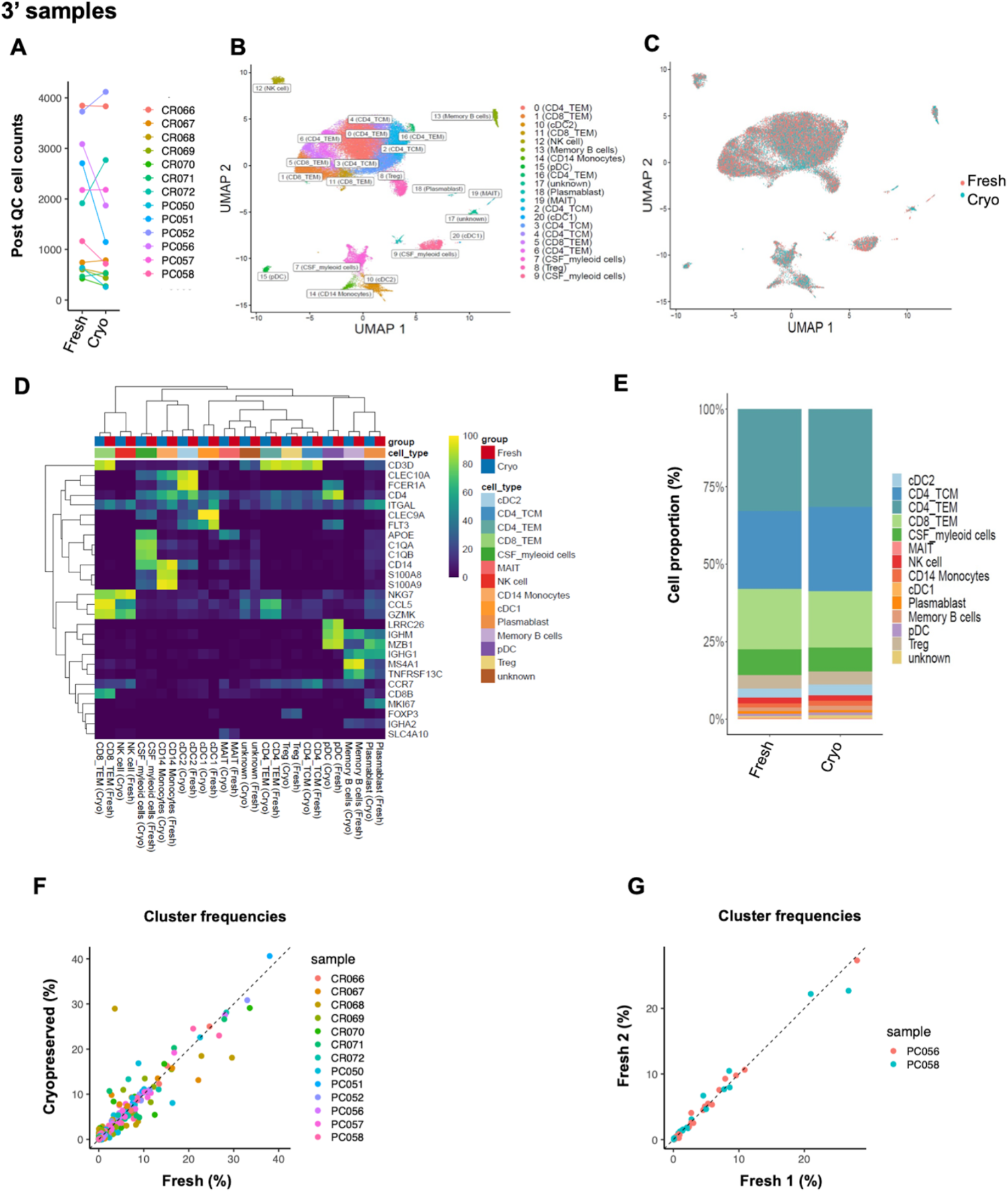
Efficient CSF-cell cryopreservation validated by 3’ single cell-transcriptomics. Post quality control (QC) CSF-cell counts in the fresh and cryopreserved (Cryo) sample pairs **(A)**. The fresh and Cryopreserved (PC050) sample pair is excluded from the differential abundance analysis due to the low number of cells (<500 cells) recovered from the cryopreserved sample. Uniform Manifold Approximation and Projection (UMAP) plot of 21 clusters/cell states color coded by their annotations (**B)**. UMAP indicating good representation of fresh and cryopreserved cells in each cluster (**C)**. Annotation of clusters using selected marker genes (**D)**. Heatmap colors correspond to the proportion of cells in each cluster expressing marker gene Bar chart indicating similar cluster proportions in five fresh and cryopreserved sample-pairs (**E)**. Significant positive correlation of each cluster proportion between five fresh and cryopreserved sample pairs (**F)**, and between two fresh-fresh sample pairs. Each point represents a cluster color coded by its sample ID (**G)**.

#### 5’ samples

None of the samples contained RBCs, and there was no significant difference in the frequency of excluded low-quality cells between fresh and cryopreserved samples (p = 0.37) (Supplementary Table 1). After excluding doublets and low-quality cells, we analyzed 24, 989 cells from 16 samples (8 pairs of fresh and cryopreserved samples) (Table 1). The number of QC pass cells were reduced in cryopreserved samples: (median=1,099 cells, IQR=825 in fresh versus median= 638, IQR=1,309 in cryopreserved, p = 0.03) (Figure 3A). However, all sequencing metrics were similar between fresh and cryopreserved samples: the percentage of mitochondrial transcripts/cell in fresh (median= 1.7%, IQR= 0.35) versus cryopreserved (median= 1.9%, IQR= 0.75) (p = 1); the number of UMI/cell in fresh (median =1,860, IQR= 757) versus cryopreserved (median=1,629, IQR= 265) (p = 0.54) ; the number of genes/cell in fresh (median = 795, IQR= 279) versus cryopreserved (median= 712, IQR= 64) (p = 0.54) (Supplementary Figure 1B). Overall, the two sets of results are consistent: Although there might be slight differences between fresh and cryopreserved samples in the number of recovered cells and the sequencing metrics, cryopreserved CSF cells generate data that meet QC parameters for downstream analyses. The cryopreservation protocol is also robust to site/batch effects and different single-cell chemistries (3’ vs. 5’) in recovering cryopreserved CSF cells.

**Figure 3:**
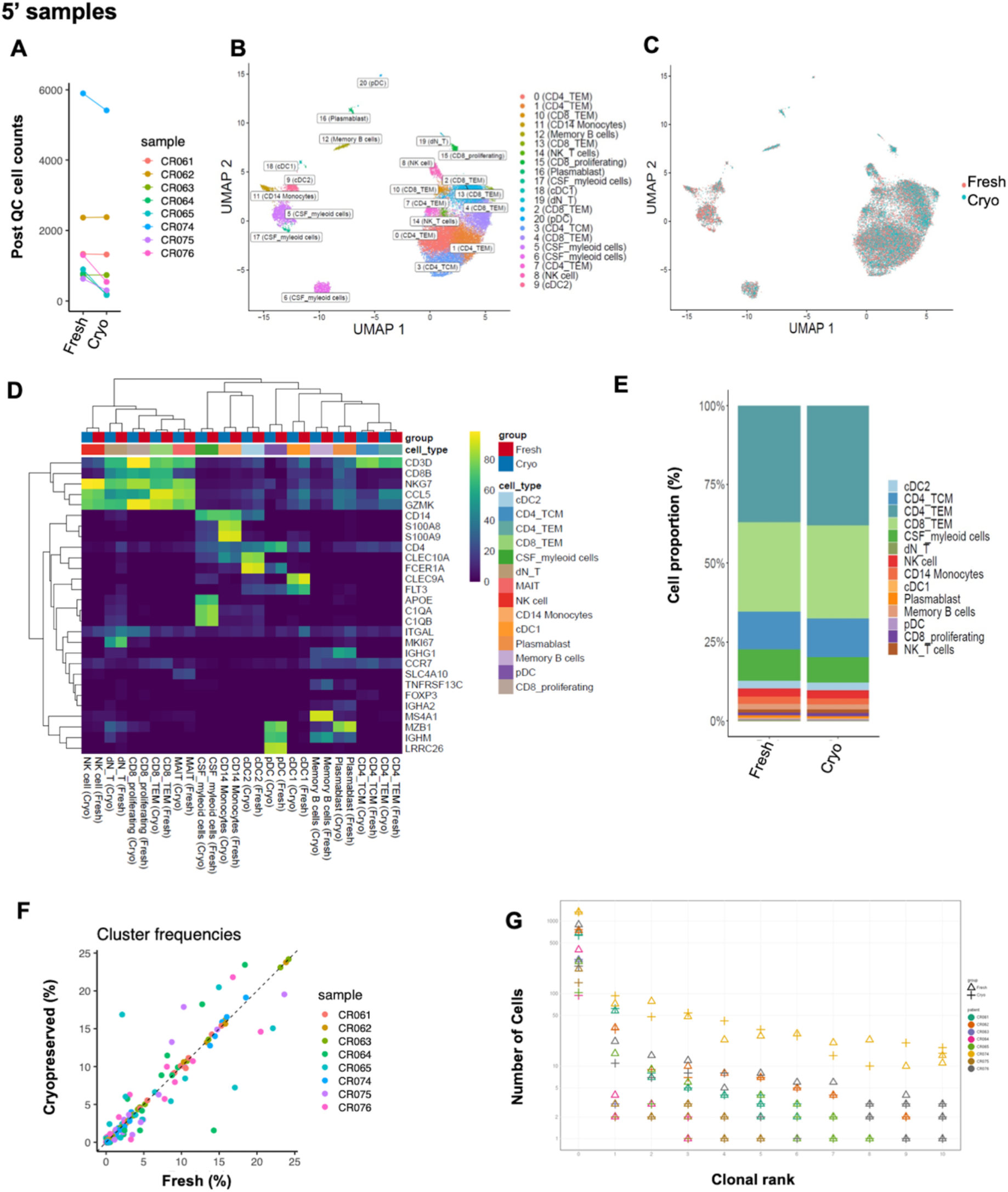
Efficient CSF-cell cryopreservation validated by 5’ single cell-transcriptomics. Post quality control (QC) CSF-cell counts in the fresh and cryopreserved (Cryo) sample pairs **(A)**. The fresh and cryopreserved (CR064 & CR065) CSF sample pairs with cryopreserved cell counts <500 cells are excluded from the differential abundance analysis. Uniform Manifold Approximation and Projection (UMAP) plot of 21 clusters color coded by their annotations (**B)** UMAP indicating good representation of fresh and cryopreserved cells in each cluster (**C)**. Annotations of clusters using selected marker genes. Heatmap colors correspond to the proportion of cells in each cluster expressing marker gene (**D)**. Bar chart indicating similar CSF-cell cluster proportions in three fresh-cryopreserved sample pairs (**E)**. Significant positive correlation of each cluster proportion in three fresh-cryopreserved sample pairs (**F)**. Clonal rank plot indicating similar distribution of the T and B-cell clonotypes in the three fresh and cryopreserved CSF samples (**G)**.

### Identification of all major CSF cell types

Normalization, integration and clustering of cells was performed independently on the 3’ and 5’ samples (see Methods for a detailed description). We identified 21 clusters of CSF cells in each dataset (Figure 2 B and Figure 3B). The top 10 marker genes for each cluster are described in Supplementary Tables 2 and 3 for 3’ and 5’ samples respectively. Prior single-cell studies of CSF have reported CSF-specific microglia-like clusters ^(7, 16, 17)^. We refined our Azimuth-based annotations for myeloid clusters using known marker genes for CSF-specific cells ^(7, 16, 17)^, and annotated clusters 7 and 9 in the 3’ samples (Figure 2B) and clusters 5, 6 and 17 in the 5’ samples (Figure 3B) as “CSF-myeloid” cells. Figure 2D and Figure 3D illustrate each cluster marker gene expression in 3’ and 5’ samples respectively. Sequencing QC metrics between fresh and cryopreserved cells showed similar patterns across individual clusters suggesting that the cryopreservation did not disproportionately affect RNA quality of any specific CSF-cell type (Supplementary Figure 2). We identified all major cell types previously reported to be present in CSF.

### Cryopreservation does not result in loss of CSF cell types

We assessed the sensitivity of CSF-cell types to cryopreservation by comparing their proportion/frequency between fresh and cryopreserved samples. The counts and percentage of cells found in each cluster from the 3’ and 5’ samples are available in (Supplementary Table 1).

We observed high correlation of cell type frequencies between all fresh-cryopreserved sample pairs (median Spearman correlation = 0.93 for 3’ samples, and 0.97 for 5’ samples) (Figure 2F, 3F and Supplementary Figure 3). Correlation was slightly higher between the two fresh-fresh replicate pairs from a subset of the 3’ samples (0.98 and 0.99) (Figure 2G). PCA of cluster frequencies further highlights that the fresh and cryopreserved cells from individual samples are more similar to each other rather than they are to their sample class (fresh or cryopreserved) (Supplementary Figure 4). However, both correlation and PCA analysis suggest that cryopreserved samples with substantially low numbers of recovered cells (e.g., CR064FRZ and CR065FRZ, both <200 cells) show higher variability in deviance from their expected cluster frequencies. Hence, we recommend that researchers exercise extra caution when analyzing data from samples with lower-than-expected recovered cell counts.

We also compared differences in each cell cluster frequency between fresh and cryopreserved sample pairs using the Wilcoxon signed-rank test. We observed no significant difference in the frequencies of each broad cell type (eg. when all CD4 sub-clusters are aggregated as one cell type) or individual cell clusters in 3’ samples (Figure 2E, Supplementary Figure 5A) and 5’ samples (Figure 3E, Supplementary Figure 5B) (all adjusted p >0.05). These results suggest that our cryopreservation method does not result in significant loss of any specific CSF cell types.

### Cryopreservation does not affect gene expression of CSF-cells

We next assessed the impact of cryopreservation on gene expression in CSF-cell types. We observed a high correlation in gene expression between fresh and cryopreserved cells across all 21 clusters in 3’ samples (median correlation = 0.99, IQR= 0.005, Supplementary Figure 6), and in 5’ samples (median correlation = 0.99, IQR= 0.01, Supplementary Figure 7). We observed a low number of differentially expressed genes between fresh and cryopreserved cells both in the 3’ samples (between 0-10 genes with fold-change >1.5 across 21 clusters, Supplementary Figure 6, Supplementary Table 4) and in the 5’ samples (between 0-4 genes with fold-change >1.5 across all 21 clusters, Supplementary Figure 7, Supplementary Table 5).

Among differentially expressed genes, hemoglobin HBB, and mitochondrial MT-ND4L gene, mitochondrial-like paralogs MTRNR2L8 and MTRNR2L12, lncRNA genes AL138963.4 and SNHG25, histone gene HIST1H1E, and PABPC1 and MYH9 genes showed reduced expression, while ribosomal gene RPS20 and mitochondrial gene MT-ATP8 showed increased expression in more than one cell type in either 3’ or 5’ cryopreserved samples (Supplementary Figure 8A, B), suggesting that cryopreservation affects these genes in a non-cell type specific manner. HBB and MTRNR2L8 were the only 2 genes that were differentially expressed in more than 1 cluster with >2 fold change. MTRNR2L8 is implicated as an antiapoptotic factor, and its decreased expression might be related to the biological effects of cryopreservation on cell survival or transcriptome. On the other hand, HBB, which encodes hemoglobin subunit beta and is highly expressed in RBCs, was mostly found in RBC-contaminated fresh samples and probably represents contamination with ambient RNA in these samples (Supplementary Figure 8 C, D). These results suggest that our cryopreservation method maintains the overall gene expression profile of CSF cell types except for very few genes.

### CSF clonotypes are conserved after cryopreservation

Lastly, we assessed the effect of cryopreservation on the frequency of T and B-cell clonotypes in 5’sc-RNAseq data. The number of cells identified from each clonotype and their corresponding CDR3 amino acid sequences are available in Supplementary Table 6. 77% of the cells in the T-cell clusters and 87 % of the cells in B or plasma cell clusters had an identified CDR3 sequence. There was no difference in the frequency of cells with identified CDR3 sequence between fresh and cryopreserved samples in the T-cell or B-cell components (p = 0.44 and 0.2, respectively). Of all identified unique CDR3 sequences, 25% were commonly found in fresh and cryopreserved samples. The non-shared clones were equally distributed between fresh and cryopreserved samples, and all had lower frequencies (Supplementary Figure 9A). The frequency of the shared clones was highly correlated between paired fresh and cryopreserved samples (Spearman’s ρ = 0.94, p <0.001, Supplementary Figure 9B). Within each patient, T and B-cell clonotypes were identified by their unique CDR3 sequence, and ranked using their occurrence frequency (i.e., clonal expansion). Figure 3G is a clonal rank plot showing that both expanded clones (rank 1 to 10) and unexpanded clones (rank 0), remain unaffected by cryopreservation (overlapping ‘triangles’ and ‘crosses’ of the same color). Altogether, these findings suggest that cryopreservation has no effect on the diversity (or lack thereof) of the adaptive immune receptor repertoire.

## Discussion

High quality peripheral blood and CSF-lymphocyte samples are imperative for elucidating the cellular and molecular cascades orchestrating neuroinflammatory and neurodegenerative disorders. Recent single-cell RNAseq investigations in fresh CSF samples have allowed comprehensive fine-grained mapping of CSF-cell types, transient cell states and disease-associated cell-type specific signatures^(17, 20, 21)^, observations that cannot be achieved using conventional flow cytometry in CSF. Cryopreservation of CSF-cells using a robust protocol that preserves cellular and molecular phenotypes would significantly advance basic and clinical research. Recently, Oh and colleagues^(22)^ published a CSF cryopreservation protocol; however, no thorough comparison of fresh and cryopreserved CSF samples was conducted. We developed a simple, yet rigorous and cost-efficient protocol to cryopreserve CSF-cells that showed highly reproducible results at two independent sites when data from fresh and cryopreserved CSF-cells are compared. We reported in an earlier study^(23)^ in human PBMCs that our cryopreservation method performs better than other cryopreservation media in recovery of immune cells with minimal impact on gene expression profile.

The CSF protocol enabled us to analyze between 165-5414 cryopreserved cells, depending on the volume of the starting sample, which was as low as 3mL, and on the participant’s diagnosis. We recovered >70% of cryopreserved cells post-thawing using our protocol at both sites. We validated the efficiency of our protocol in recovering good quality cells using two single cell RNA-sequencing chemistries, the 3’ sc-RNA-seq and the 5’ sc-RNA-seq, which are known to have different sensitivity to detect genes. We identified all known CSF-cell types including microglia-like cells reported by others that we report as “CSF myeloid-cells” given the lack of clarity as to their origin. Our cryopreservation method did not alter the composition of CSF-cells; the frequency and gene expression profiles of cell types (even minor cell subsets like B-cells) were highly preserved. We noted a higher hemoglobin gene expression in the fresh samples despite the exclusion of RBCs from the analyses. However, no RBCs were found in our cryopreserved pairs for RBC-contaminated fresh samples suggesting that the cryopreservation protocol might be beneficial in removing RBC contamination and hemoglobin ambient RNA, when present. Finally, our cryopreservation protocol is able to preserve T and B-cell clonotypes.

## Conclusion

We developed an efficient and reliable cryopreservation method for long-term storage of CSF-cells, validated by high-resolution single cell analysis. Our findings highlight comparable fresh and cryopreserved CSF-cell profiles. CSF-cell clusters and T/B-cell clonotypes in fresh *versus* cryopreserved samples are preserved and comparable. It is possible that some rare and functionally relevant immune subtypes may not be recovered after cryopreservation, a possibility which should be explored in future studies using multimodal assays such as CITE-seq^(24)^. Given the practical challenges of fresh CSF characterization in large multi-center settings with different levels of technological sophistication and considering cost savings, adopting the proposed method will enable large-scale and multicenter investigations - crucial for clinical trials.

## Data availability

Seurat objects containing gene expression and metadata from cells included in the analyses from the thirteen pairs of 3’ samples and eight pairs of 5’ samples are available as Supplementary Data Files 1 and 2. The gene expression data from Basel and Columbia samples are available via an interactive browser https://malhotralab.shinyapps.io/csf_cryo_3prime/ https://malhotralab.shinyapps.io/csf_cryo_5prime/

## Contributions

DM, PLD and HT designed the study. HT and DC carried out experiments. TR analyzed data. HT, TR, and DM drafted the manuscript. DC, SE, IC, TD performed the lumbar punctures. CR and LC contributed to the study design, preparation of protocol video and interpretation of results. VM, DM, PDJ, TR and HT contributed to the analysis plan and interpretation of results. DM and PDJ conceived and supervised the overall project and edited the manuscript.

## Acknowledgments

We thank all patients who generously offered CSF and blood samples. We are grateful to the single cell sequencing core of Columbia University and FGCZ at the University of Zurich. We thank Imran Fanaswala in the DM lab for help with the figures. We are grateful to the entire Roche Consonance Team for their help with the implementation of the CSF cryopreservation protocol and sharing the video.

Video link: https://mf.wetransfer.com/downloads/e93e60af1ef6cfa9a94cff342af5258420210104135502/259ac3

---

**Supplementary Figure 1:**
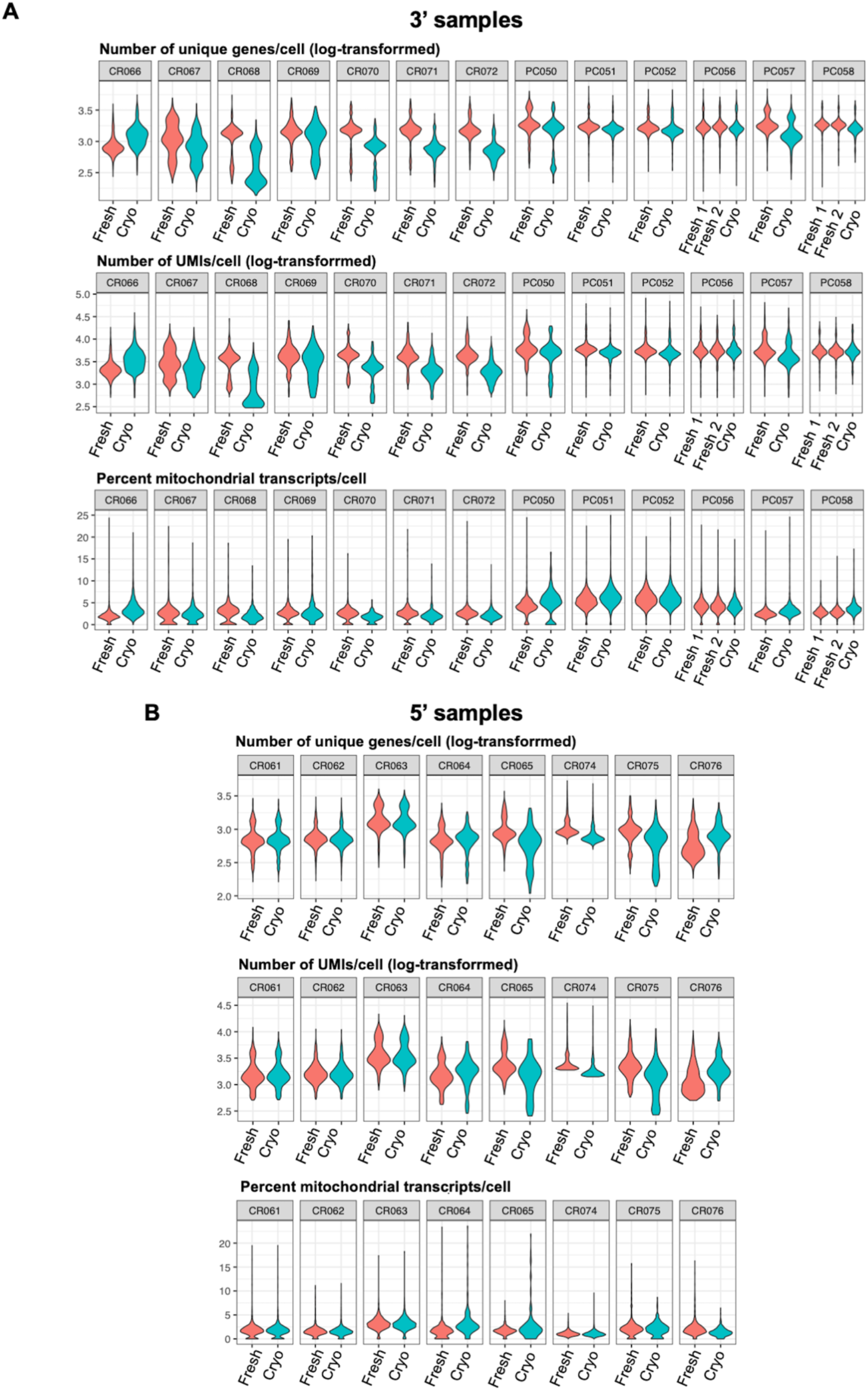
Sample level RNA-sequencing quality control (QC). Sequencing QC from 26 samples sequenced using 3’ chemistry across Columbia university and University Hospital Basel **(A)**, and 16 samples using 5’ sequencing chemistry across Columbia university and University hospital Basel samples **(B)**. Violin plot of the number of unique genes **(A-B)**, number of unique transcripts **(A-B)**, and percentage of mitochondrial reads **(A-B)** in each fresh-cryopreserved sample-pair. The values of number of detected genes and transcripts is on the log scale.

**Supplementary figure 2:**
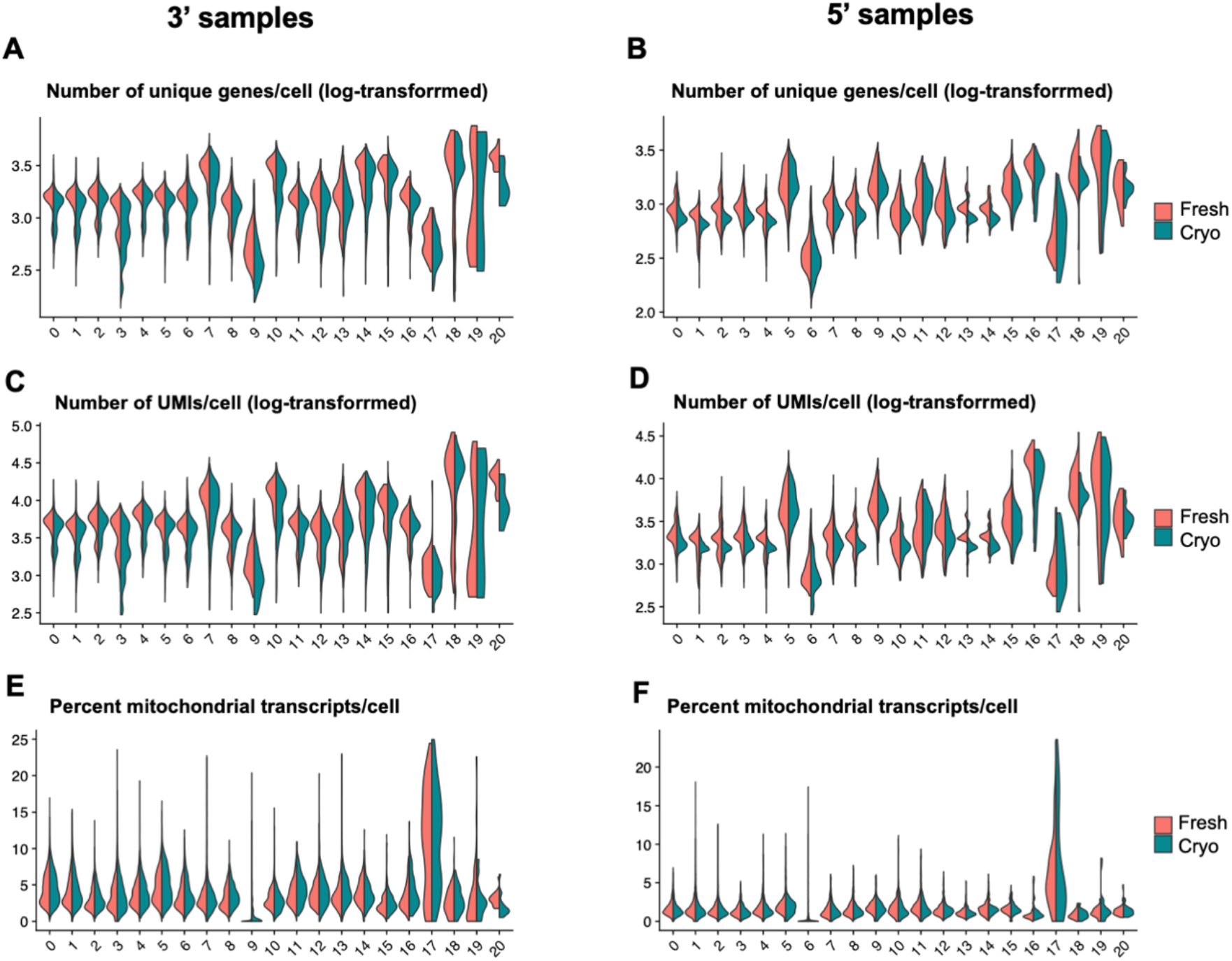
Cell cluster level RNA sequencing quality control (QC). Sequencing QC in 21 clusters from 3’ samples **(A-C-E)**, and in 21 clusters from 5’ samples **(B-D-F)**. Density plot showing the similar distribution of the number of genes **(A-B)**, number of unique transcripts **(C-D)**, and percentage of mitochondrial genes in each cluster **(E-F)**. The values of number of detected genes and transcripts is on the log scale. Pink colored density plots represent fresh cells and blue colored plots represent cryopreserved cells in each cluster.

**Supplementary Figure 3:**
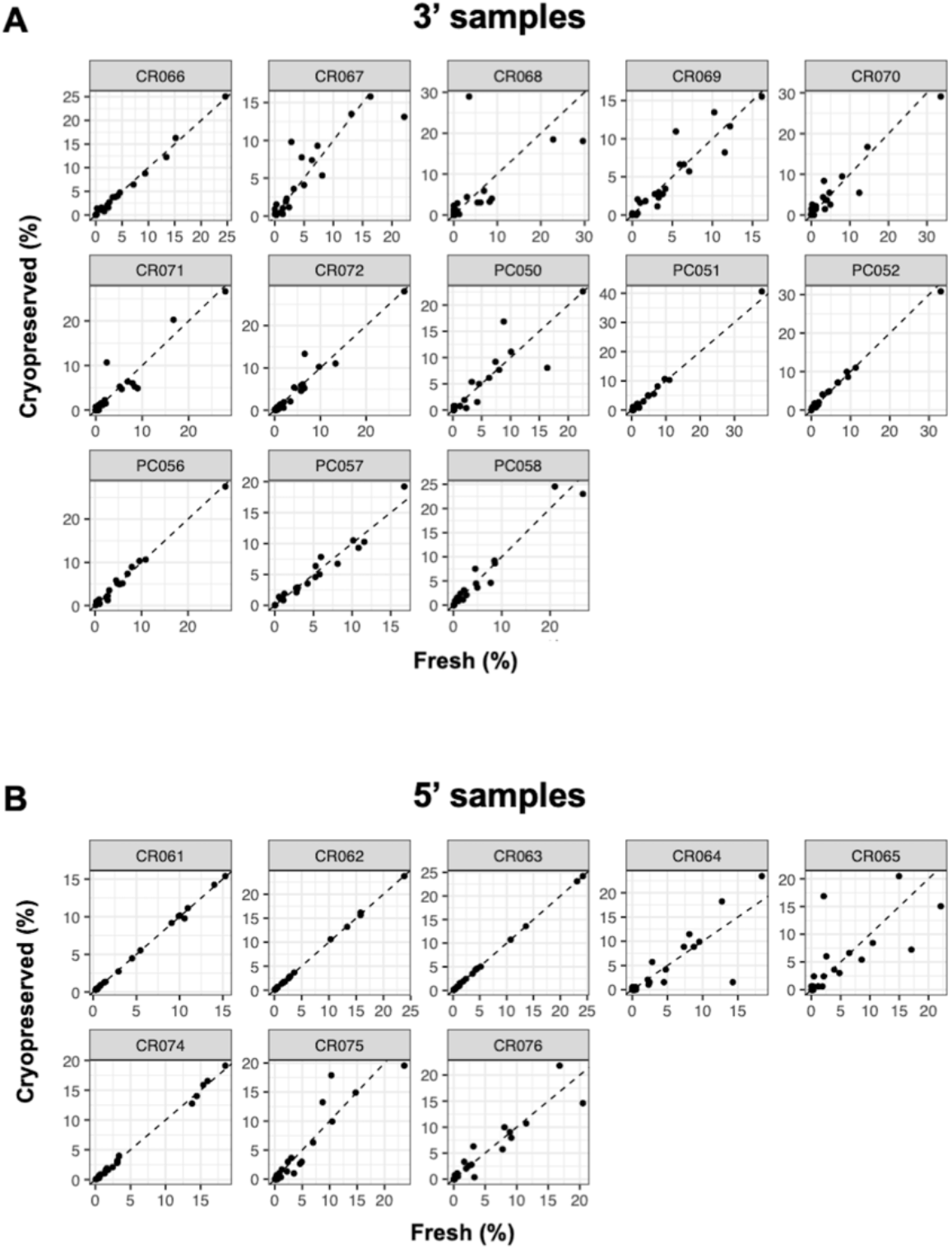
Correlation of cluster distribution (%) within fresh or cryopreserved samples, analyzed using 3’ **(A)** and 5’ **(B)** sequencing chemistry, respectively.

**Supplementary figure 4:**
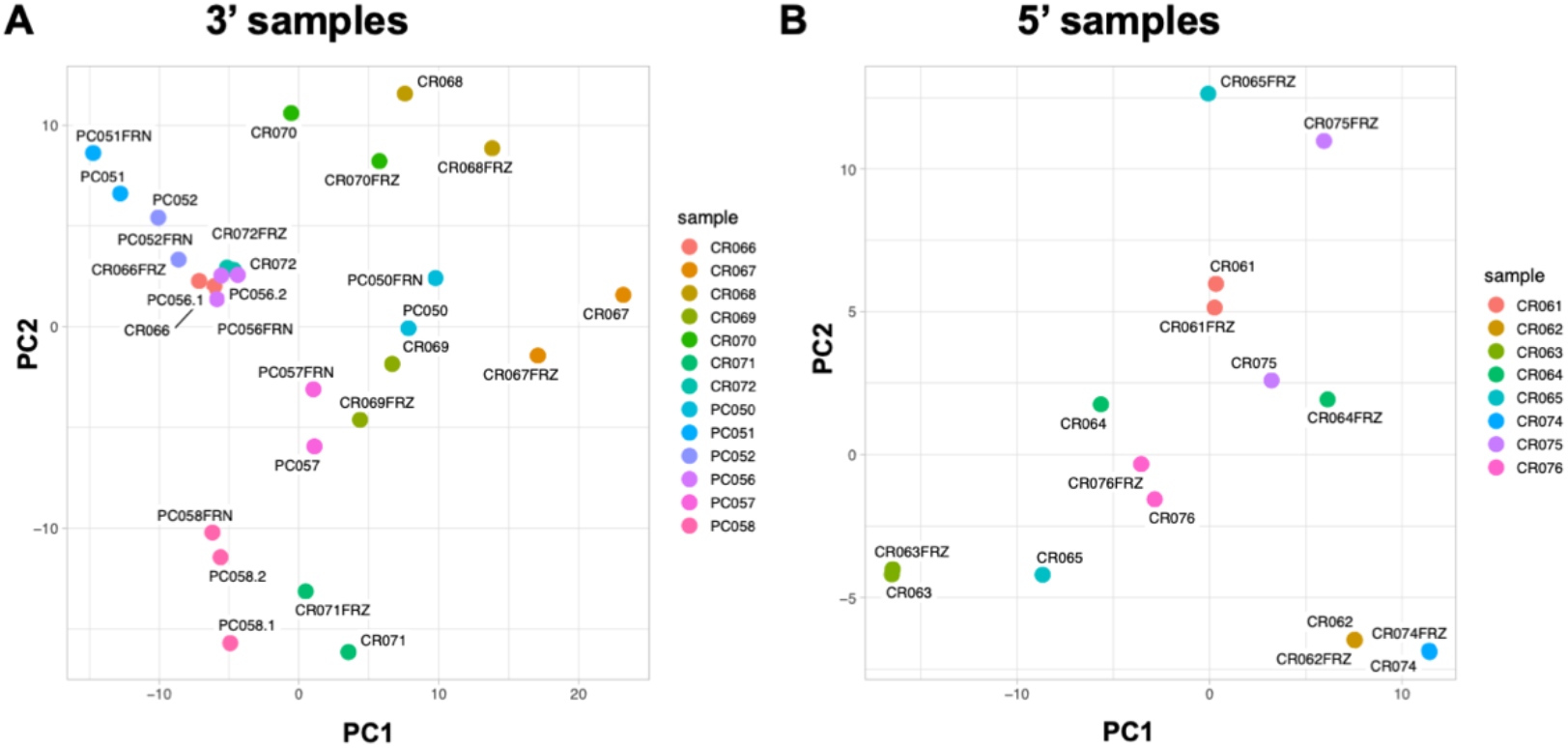
Principal Component Analysis (PCA) clustering of samples based on their cluster frequencies. **(A)** 3’ sequencing chemistry samples, **(B)** 5’ sequencing chemistry samples.

**Supplementary Figure 5:**
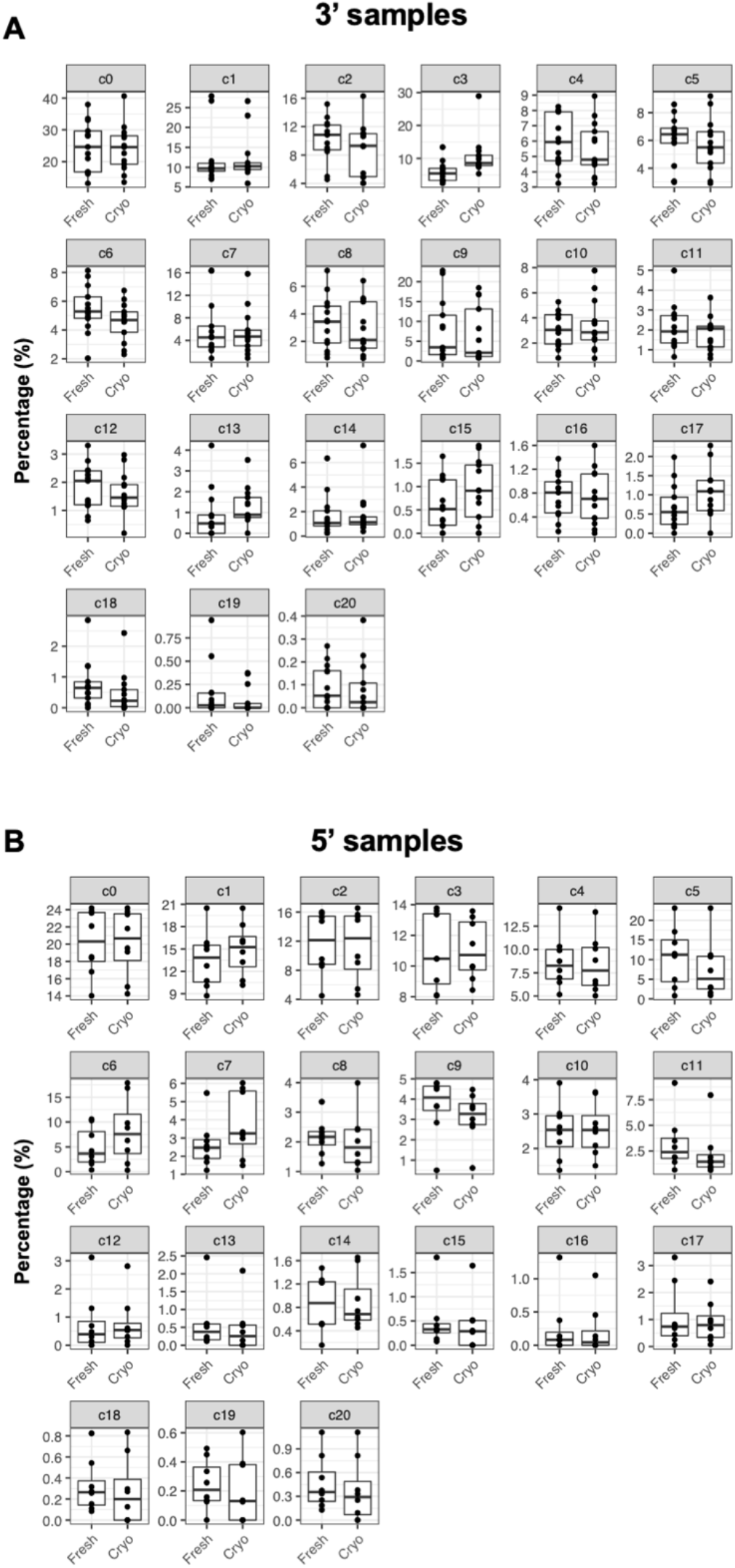
Frequencies of individual clusters in fresh and cryopreserved samples sequenced using 3’ **(A)**, and 5’ **(B)** sequencing chemistry, respectively.

**Supplementary Figure 6:**
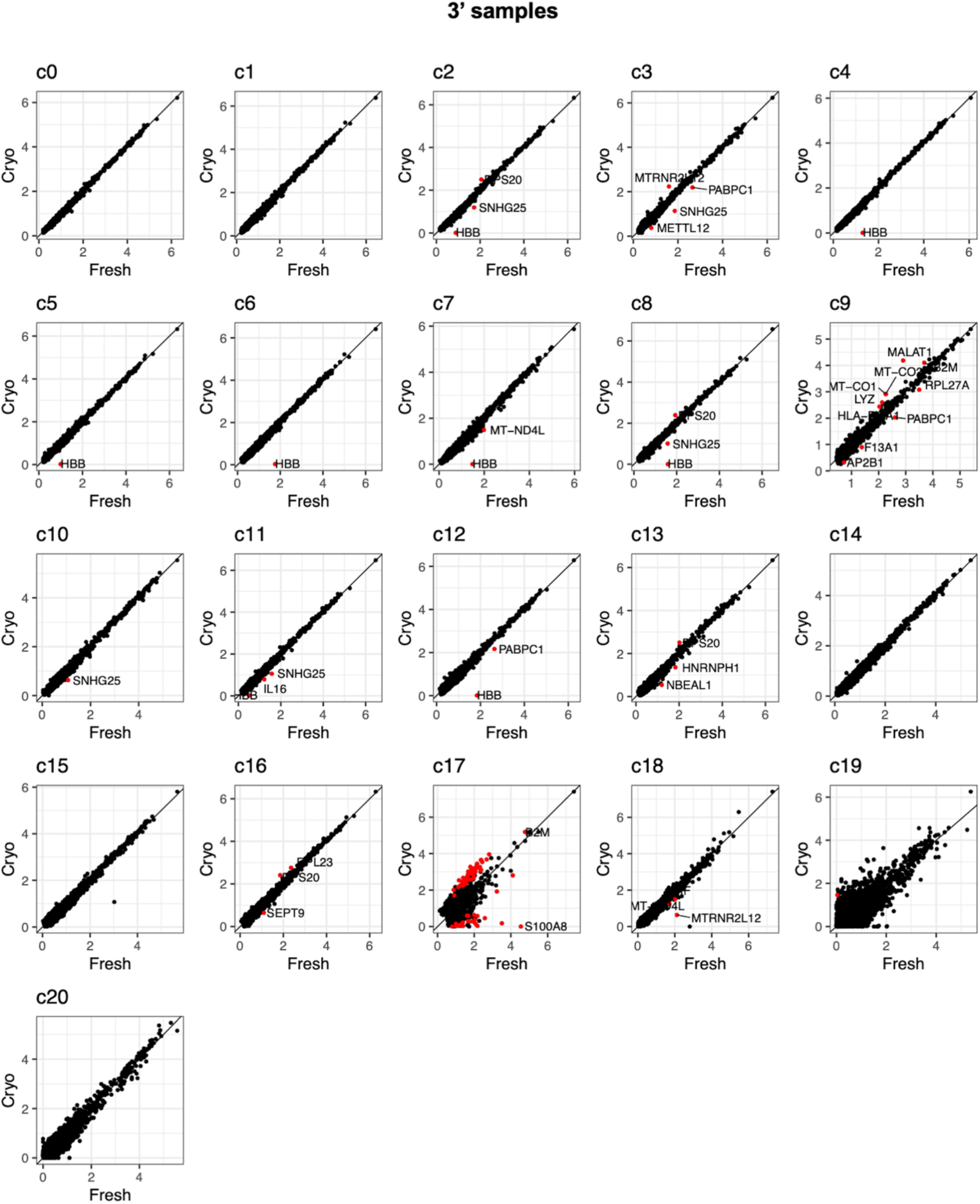
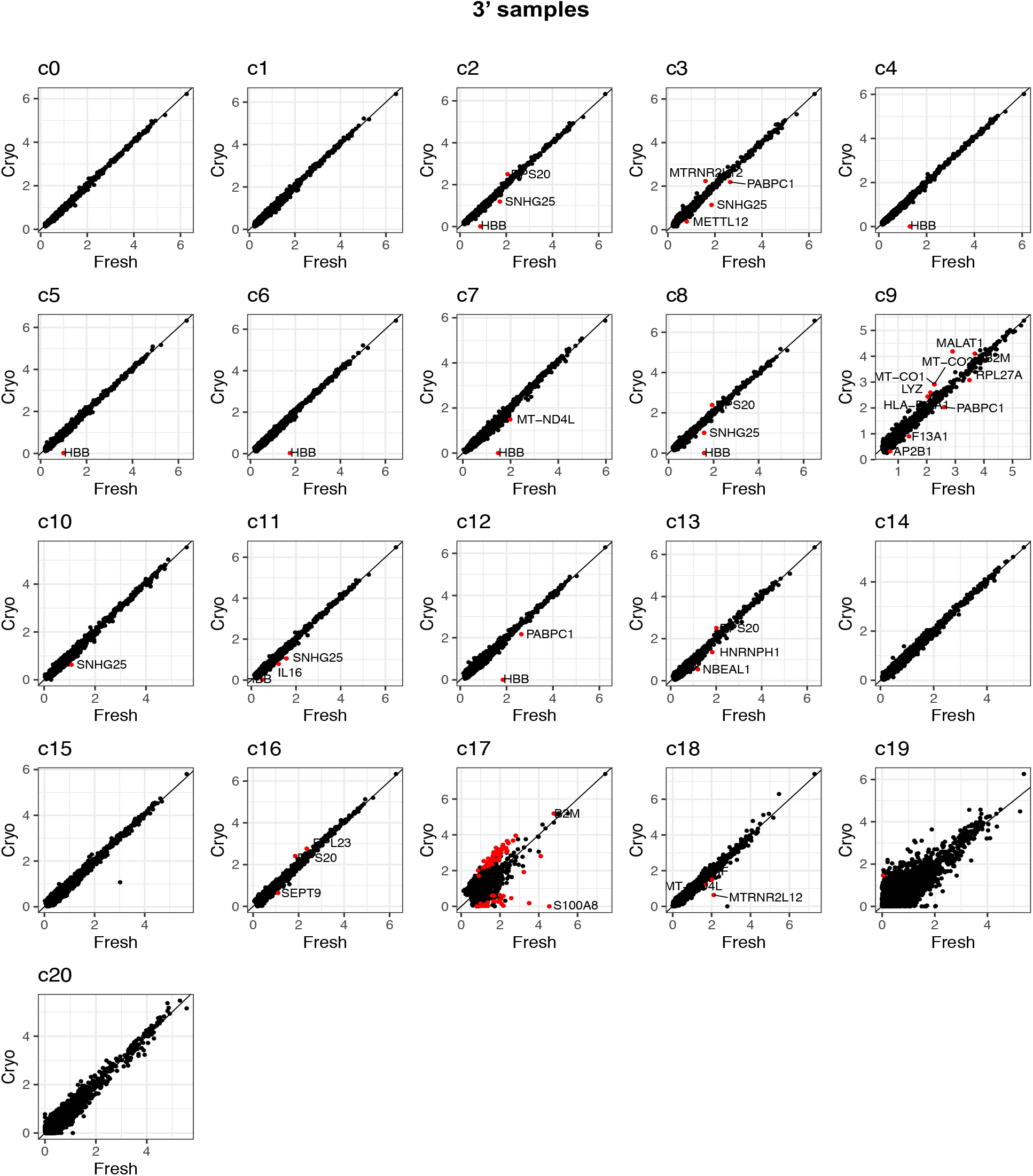
Gene expression correlations in each cluster in fresh compared to cryopreserved CSF samples, processed using 3’ sequencing chemistry. Differentially expressed genes are labeled and shown in red.

**Supplementary Figure 7:**
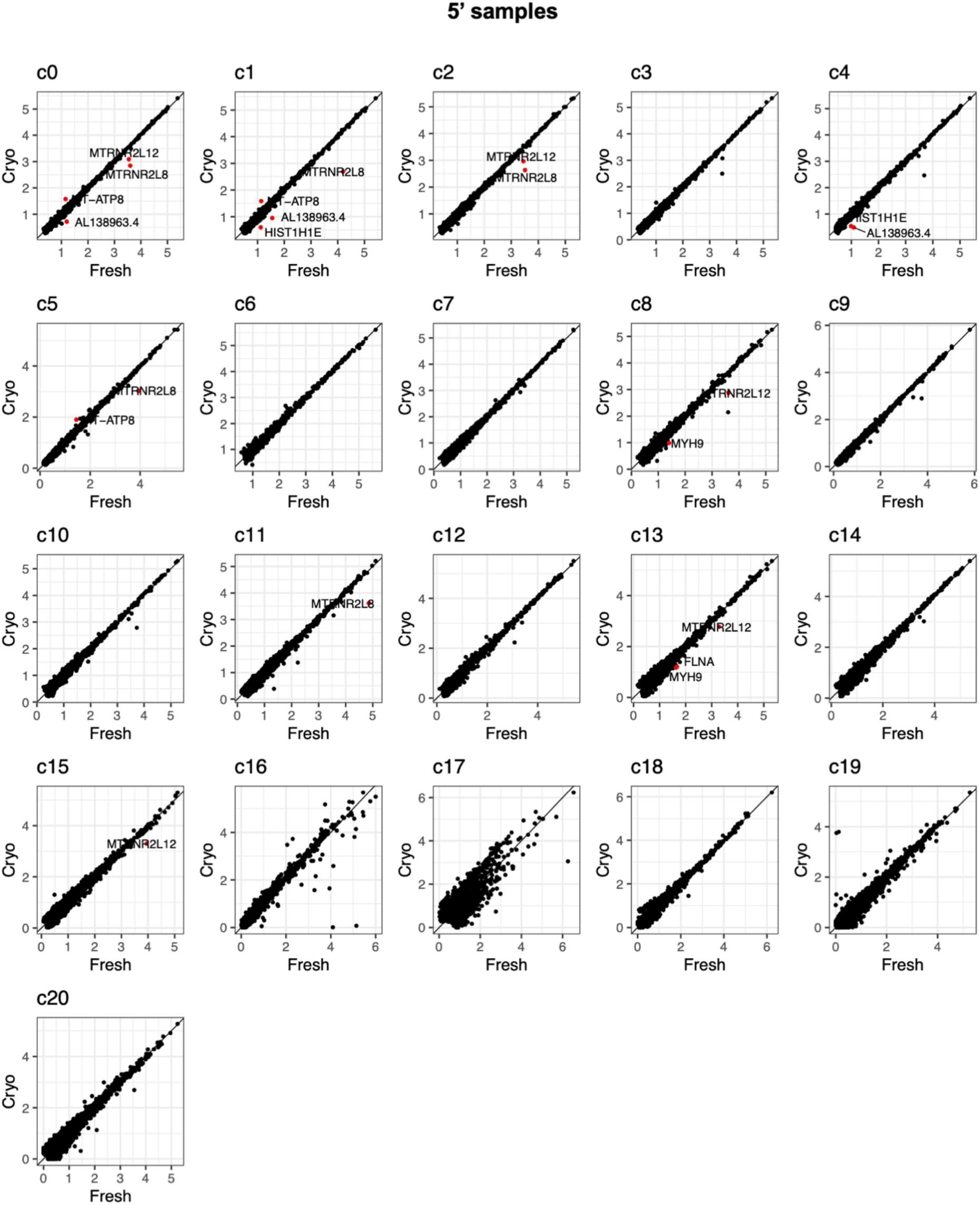
Gene expression correlations in each cluster in fresh compared to cryopreserved CSF samples, processed using 5’ sequencing chemistry. Differentially expressed genes are labeled and shown in red.

**Supplementary Figure 8:**
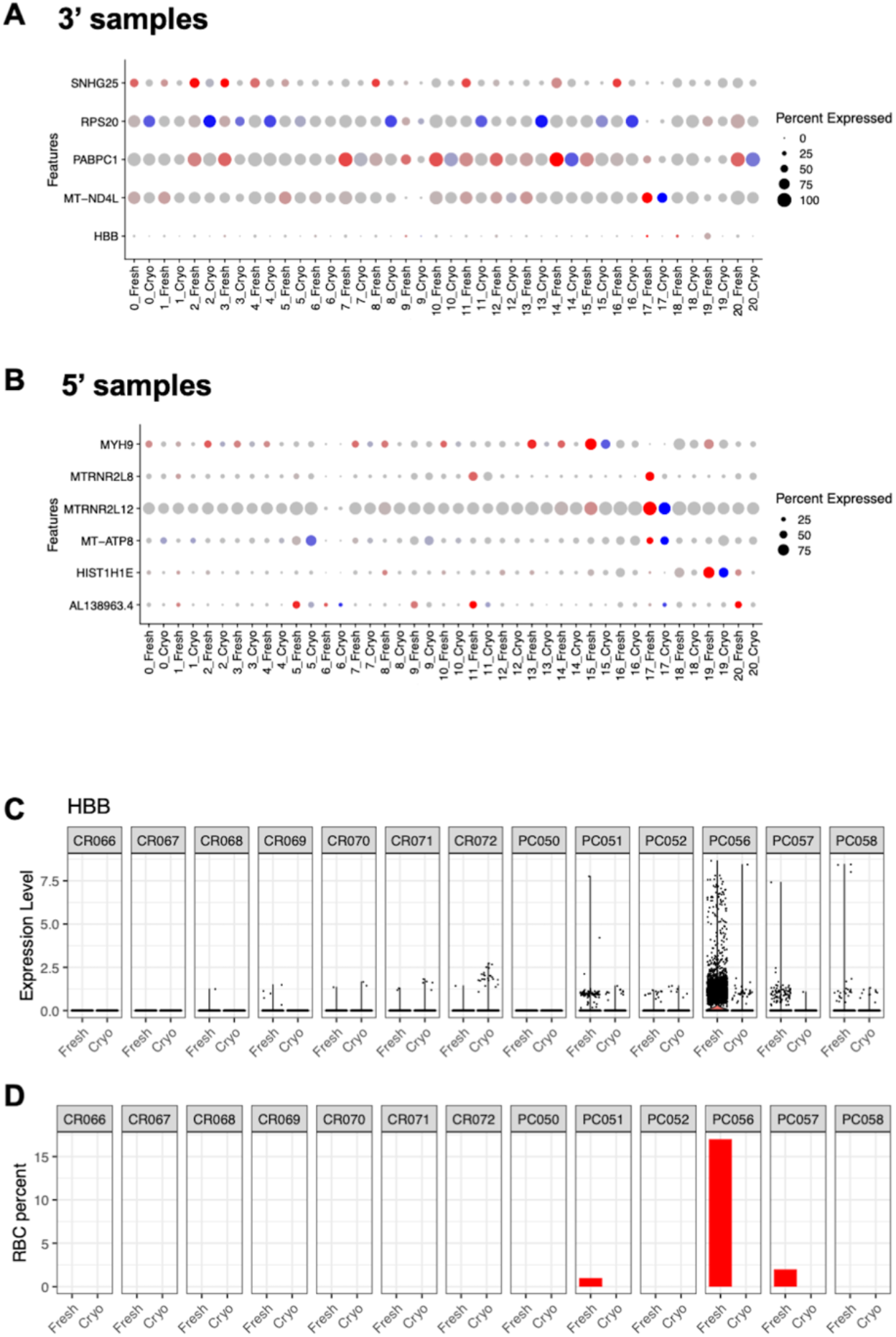
Genes susceptible to change in response to the cryopreservation process in 3’ sequencing chemistry samples. **(A)** and **(B)** 5’ sequencing chemistry samples **(C)** The normalized expression levels of *HBB* in cells in five sample-pairs. The percentage of red blood cells identified in each sample before exclusion **(C)**. Genes with higher expression in clusters from fresh (red dots) and cryopreserved samples (blue dots) **(D)**. Darker color represents higher expression. Dot size represents the percentage of cells in each cluster in which the gene is expressed.

**Supplementary Figure 9:**
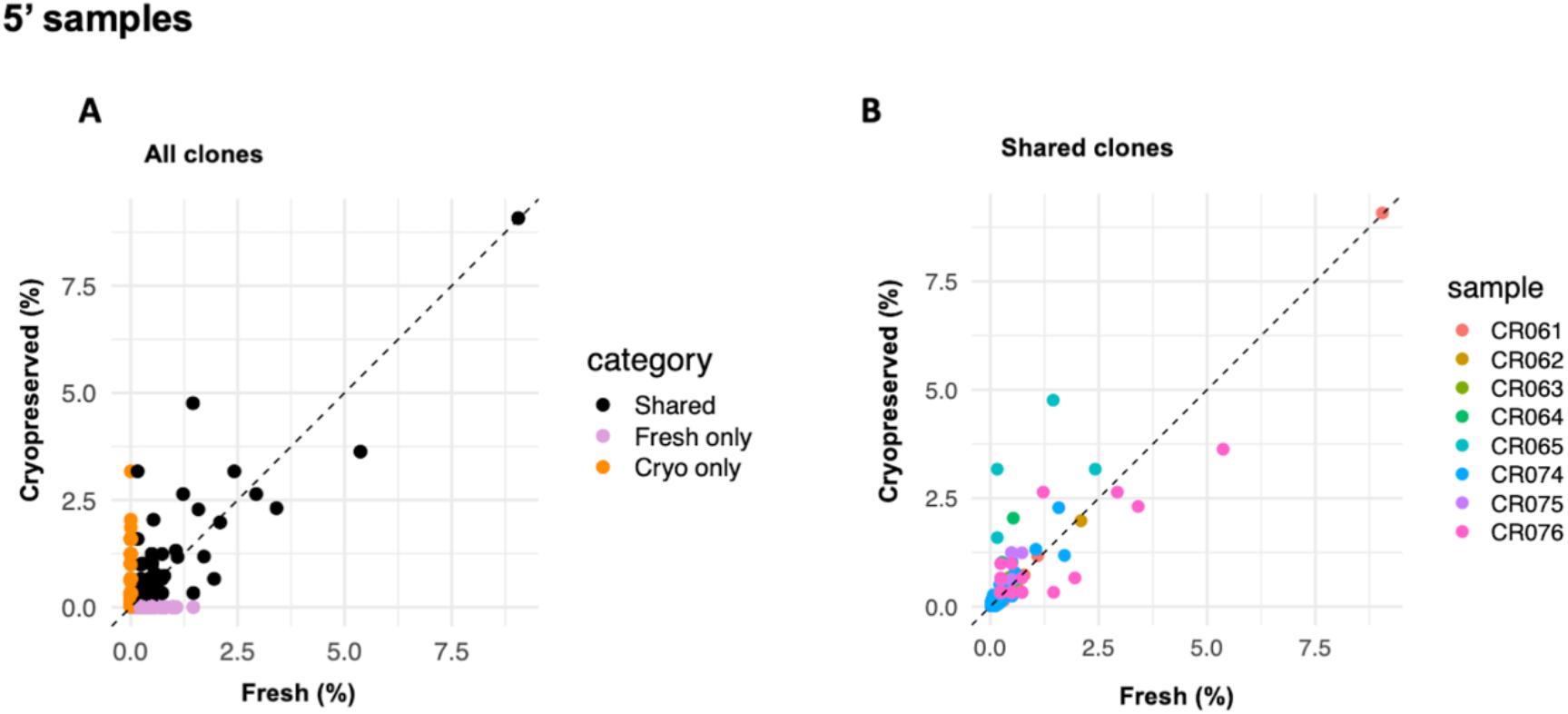
Clonotype sharing between fresh and cryopreserved CSF samples and across each cluster processed using 5’ sequencing chemistry. Clonotype percentage of frequencies **(A)**. Correlation between clonotype frequencies shared between fresh and cryopreserved sample pairs **(B)**.

